# Bottlebrush polyethylene glycol nanocarriers translocate across human airway epithelium via molecular architecture enhanced endocytosis

**DOI:** 10.1101/2024.02.15.580508

**Authors:** Zhi-Jian He, Baiqiang Huang, Li-Heng Cai

## Abstract

Pulmonary drug delivery is critical to the treatment of respiratory diseases. However, the human airway surface presents multiscale barriers to efficient drug delivery. Here we report a bottlebrush polyethylene glycol (PEG-BB) nanocarrier that can translocate across all barriers within the human airway surface. Guided by the molecular theory, we design a PEG-BB molecule consisting of a linear backbone densely grafted by many (∼1,000) low molecular weight (∼1000 g/mol) PEG chains; this results in a highly anisotropic, wormlike nanocarrier featuring a contour length of ∼250 nm, a cross-section of ∼20 nm, and a hydrodynamic diameter of ∼40 nm. Using the classic air-liquid-interface culture system to recapitulate essential biological features of the human airway surface, we show that PEG-BB rapidly penetrates through endogenous airway mucus and periciliary brush layer (mesh size of 20-40 nm) to be internalized by cells across the whole epithelium. By quantifying the cellular uptake of polymeric carriers of various molecular architectures and manipulating cell proliferation and endocytosis pathways, we show that the translocation of PEG-BB across the epithelium is driven by bottlebrush architecture enhanced endocytosis. Our results demonstrate that large, wormlike bottlebrush PEG polymers, if properly designed, can be used as a novel carrier for pulmonary and mucosal drug delivery.

**Table of Contents:** 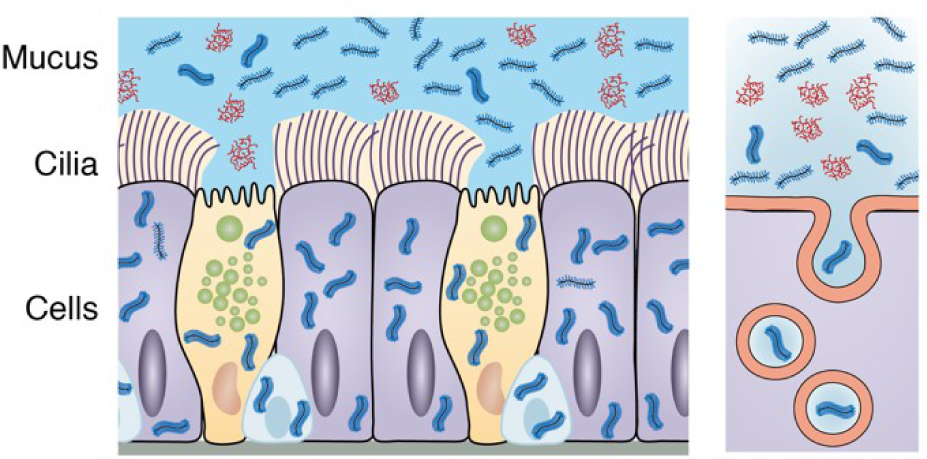

## Main Text

Pulmonary drug delivery^1^ is critical to the treatment of respiratory diseases, such as asthma^2^, chronic obstructive pulmonary disease (COPD)^3^, and pulmonary fibrosis^4,5^. However, the human airway surface has a multiple-layer structure^6–8^ that presents multiscale barriers to efficient and localized drug delivery. Lining the airway surface is mucus, a viscoelastic and sticky hydrogel that traps essentially any inhaled particulates and pathogens^9–11^. The mucus is further separated from the epithelium by a periciliary layer, which provides a favorable environment for cilia beating and cell surface lubrication (**Fig. 1A**). Together with trapped objects, the mucus hydrogel is transported out of the lung by coordinated cilia beating. While essential to maintaining respiratory health, this mucociliary clearance also prevents the retention of drugs within the airway^12,13^. Furthermore, the epithelial cells are connected by cell junctions such as tight junctions, forming an integrated epithelial barrier that prevents drug carriers from traversing and reaching the underlying cells^14–16^. Thus, efficient pulmonary drug delivery requires carriers that can quickly penetrate mucus faster than its turnover rate, sneak through the periciliary layer, and then be internalized into and retained within epithelial cells.

**Figure 1.**
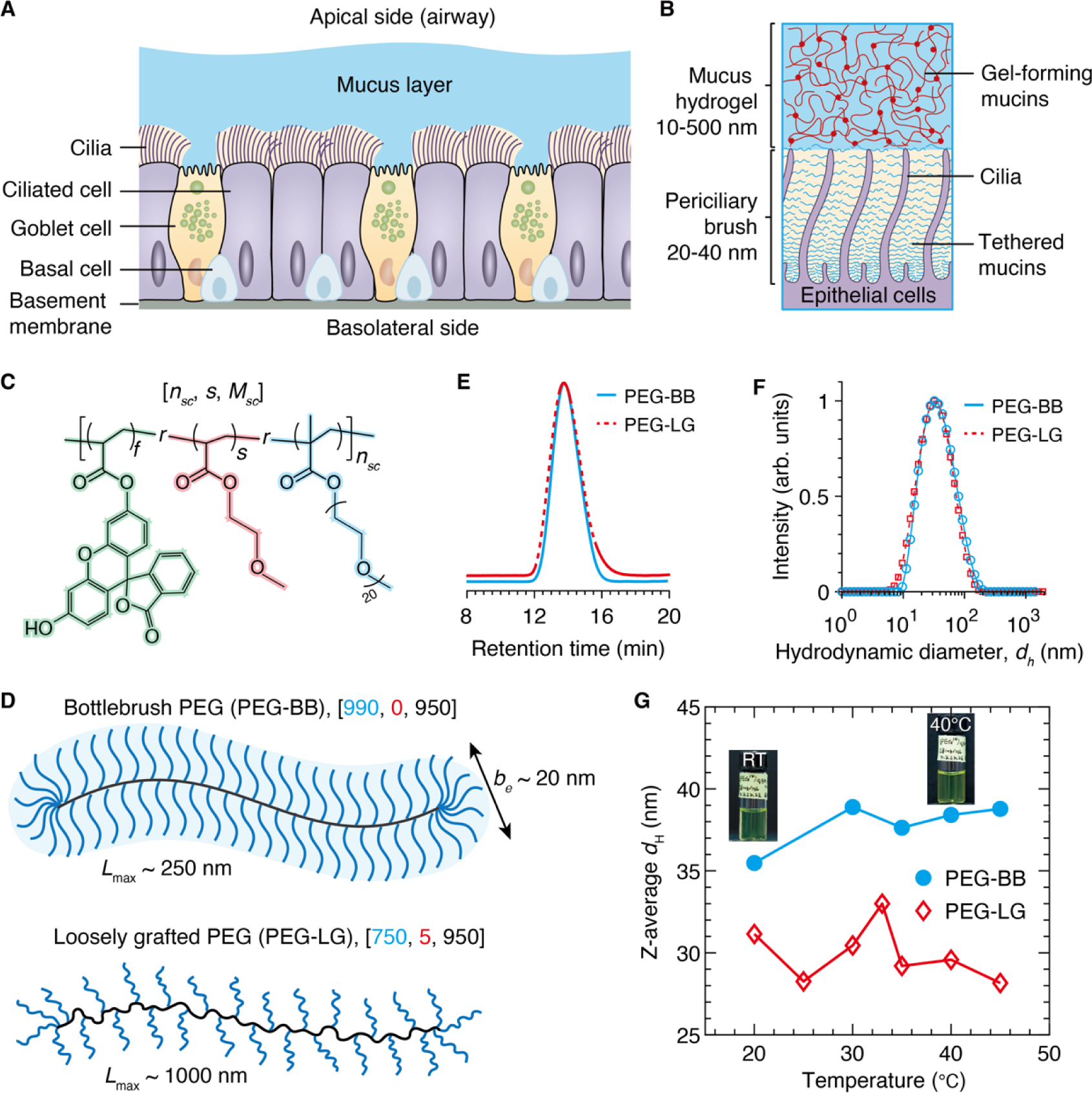
Design and synthesis of grafted PEG polymeric nanocarriers for pulmonary delivery. (A) The human airway epithelial barrier consists of three distinct layers: (i) mucus hydrogel, (ii) periciliary layer, and (iii) a pseudostratified epithelium in which the cells are connected by cell junctions. **(B)** The mucus hydrogel has a mesh size in the range of 10-500 nm. In the periciliary layer, transmembrane mucins are tethered to cilia and the epithelial surface to form a brush-like layer with a mesh size in the range of 20- 40 nm. **(C)** The grafted PEG macromolecule is a random copolymer consisting of side chains (PEG methacrylate of 950 g/mol), spacers (2-methoxyethyl acrylate), and fluorescent labels (fluorescein *O*- methacrylate). The design parameter space is [*n_sc_*, *s*, *M_sc_*], in which *n_sc_* is the number of side chains, *s* is the spacer/side chain ratio, and *M_s_* is the side chain MW. For all PEG-based carriers, the molar fraction of fluorescein, *f*, is fixed at 0.01. **(D)** Upper: A densely grafted bottlebrush PEG (PEG-BB) with 950 side chains and no spacers. The polymer is a worm-like molecule with a cross-section size of ∼20 nm and a contour length of ∼250 nm. Lower: A loosely grafted PEG (PEG-LG) with approximately 750 side chains, in which two neighboring side chains are separated by 5 spacer monomers on average. The polymer is a comb-like molecule with a cross-section size of ∼6 nm and a contour length of ∼1000 nm (**Supporting Text**). **(E)** GPC profiles of PEG-BB and PEG-LG nearly overlap, indicating the two molecules have similar hydrodynamic diameters. The polydispersity index for both polymers is the same, 1.45. **(F)** Dynamic light scattering (DLS) of PEG-BB and PEG-LG at room temperature. **(G)** The *Z*-average hydrodynamic diameter is 37 nm for PEG-BB and 32 nm for PEG-LG. Moreover, their sizes are nearly independent of temperature from 20 °C to 45 °C. The error bar is smaller than the symbol size (*n*=3).

Recent advances in drug delivery have demonstrated that coating sub-micrometer latex particles with bioinert polymers such as polyethylene glycol (PEG) promotes particle penetration through mucus^17–19^. However, as demonstrated by our previous work^6,20,21^ and others^8,9,22–25^, the mucus hydrogel is highly heterogeneous with a wide distribution in the network mesh size from tens to hundreds of nanometers; thus, the PEGylated solid particles can still be physically trapped by local small meshes. Decreasing the particle size to ∼100nm or less helps circumvent the physical confinement. However, such small nanoparticles have a large surface curvature that inevitably causes unreliable coating, such that the particles adhere to mucus through non-specific biochemical binding^26^, a phenomenon commonly known as mucoadhesion^27^. In addition, we recently discovered that the periciliary layer is not simply filled with low-viscosity physiological liquid; instead, it is gel-like with transmembrane mucins densely grafted to cilia and the cell surface, forming a brush-like gel with a mesh size of 20-40 nm^6^ (**Fig. 1B**). This periciliary brush gel serves as a protective layer additional to mucus to prevent external objects from reaching the cell surface. It is highly desired a delivery system that sneaks through not only the physical and biochemical barriers of mucus but also the tight periciliary brush gel to be efficiently internalized by epithelial cells.

Here, we report a wormlike PEG-based polymeric nanocarrier that can rapidly translocate across all barriers within the human airway surface. Guided by the molecular theory for the structure of bottlebrush polymers in good solvent, we design and synthesize a nanocarrier consisting of a long linear backbone densely grafted by many (∼1,000) low molecular weight (MW) PEG side chains (∼1,000 g/mol). This results in a bottlebrush PEG (PEG-BB) macromolecule featuring a contour length of ∼250 nm, a cross-section of ∼20 nm, and a hydrodynamic diameter of ∼40 nm. Using the classic human bronchial epithelial cell (HBEC) culture as a model system, we show that PEG-BB can rapidly penetrate through endogenous airway mucus and the periciliary brush layer to be internalized by epithelial cells across the whole epithelium. By quantifying the cellular uptake of polymeric carriers of various molecular architectures and manipulating cell proliferation and endocytosis pathways, we show that translocation of PEG-BB across the epithelium is driven by bottlebrush architecture enhanced endocytosis. Our results demonstrate that large, wormlike PEG-BB polymers, if properly designed, can be used as a novel carrier for pulmonary and mucosal drug delivery.

## Results

### Design and synthesis of PEG-based nanocarriers with different molecular architectures

The design of our nanocarrier is inspired by the molecular structure of mucins, featuring a large polypeptide backbone that is heavily glycosylated with many sugar chains^28–30^. We seek to design a mucin-like PEG-based polymer, which consists of a long linear backbone densely grafted by many relatively short PEG side chains. Because the side chains highly overlap with each other, the only way for them to avoid molecular crowding is to extend radially away from the backbone, forming a wormlike bottlebrush structure, as illustrated in **Fig. 1C** and **Fig. S1**. We hypothesize that the grafting density of PEG side chains can be precisely controlled to enable a non-sticky carrier, whereas the flexibility and wormlike geometry of the bottlebrush carrier allow it to sneak through the tight mesh of mucus and periciliary gels to be internalized by epithelial cells.

To test this hypothesis, we design a PEG-BB nanocarrier with a precisely controlled molecular architecture, denoted by three parameters, [*n*_*sc*_, *s*, *M*_*sc*_], where *n*_*sc*_is the number of PEG side chains, *s* is the molar ratio of spacer monomers to the side chains, and *M*_*sc*_ is the MW of the PEG side chain (**Fig. 1C**). To guide the design, we develop a scaling theory for the molecular structure of a bottlebrush polymer in good solvent (**Supporting Text**). Within a bottlebrush polymer, the size of the side chain is [eq. (S6)]:

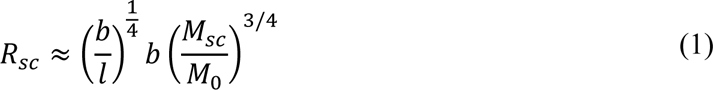

where *b* and *M*_0_ are, respectively, the Kuhn length and mass of the side chain, and *l* is distance between two neighboring grafting sites. Note that the side chain size not only increases with the grafting density 1/*l* but also scales with the polymer MW by a power of 3/4, higher than 3/5 for an unperturbed linear chain in good solvent^31^.

Analogous to “sausage versus spaghetti,” the bottlebrush polymer is essentially a wormlike ‘fat’ linear polymer^32–36^. The cross-section, or effective monomer size, of the wormlike bottlebrush is about twice the side chain size, *b*_*e*_ ≈ 2*R*_*sc*_. The radius of gyration *R*_*g*_ of the ‘fat’ linear polymer is proportional to the self-avoiding random walk of the effective monomers by a factor of α [eq. (S7)]:

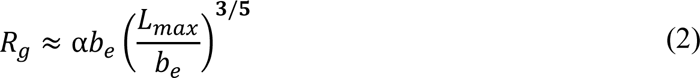

where α = 0.4 for a linear polymer in good solvent^37^ and *L*_*max*_ = *n*_*sc*_*l* is the contour length of the bottlebrush backbone.

In our experiments, we use methacrylate-terminated PEG with *M*_*sc*_=950 g/mol as the side chain. This MW is nearly the same as that used for mucus-penetrating PEGylated nanoparticles^38^. We use 2-methoxyethyl acrylate (MEA) as the spacer monomer, as MEA is chemically similar to PEG but has a much lower MW of 130 g/mol, so that it does not alter the chemical nature of the PEG carrier and only reduces the grafting density of PEG side chains. Furthermore, we use fluorescein *O*-methacrylate as the fluorescent probe and fix its fraction at 1% to ensure relatively bright fluorescence.

Using reversible addition fragmentation chain transfer radical polymerization (RAFT)^39^, a living polymerization technique widely used for controlled polymer synthesis, we copolymerize the side chain, spacer monomer, and fluorescent probe at prescribed ratios to create fluorescent PEG-based nanocarriers (**Fig. 1C**). The PEG-BB consists of 990 side chains but no spacers, [990, 0, 950], as illustrated by the upper panel of **Fig. 1D**. Successful synthesis is confirmed by proton nuclear magnetic resonance (^1^H-NMR) spectroscopy (**Fig. S2**) and gel permeation chromatography (GPC) (solid blue line, **Fig. 1E**). The linear grafting density of PEG side chains is very high, with four side chains per nanometer or *l* = 0.254 nm. Considering that for PEG in water, *b* = 0.8 nm and *M*_0_ = 44 g/mol,^40–42^ the cross-section of the PEG-BB, *b*_*e*_ ≈ 2*R*_*sc*_ ≈20 nm [eq. (1)]; this value is comparable to the lower limit of the mesh size of the periciliary brush layer. By contrast, PEG-BB has a contour length *L*_*max*_ ≈ 250 nm and *R*_*g*_ ≈ 36 nm [eq. (2)], comparable to the upper limit of the periciliary brush mesh size. Moreover, the predicted *R*_*g*_is comparable to the experimentally measured hydrodynamic diameter of PEG-BB, *d*_*h*_ = 37 nm, as shown by the dynamic light scattering (DLS) profile in **Fig. 1F** (**Supporting Text**). Further, we verify that the size of PEG-BB is nearly independent of temperature within the range between 20 °C and 45 °C (**Fig. 1G**). Because the PEG-BB has a radius of gryration that is large enough to be excluded from the periciliary brush, yet it has a cross-section smaller than the average mesh size of the periciliary brush gel, this polymer allows us to test whether the wormlike geometry allows the translocation of the PEG-BB across the human airway surface barriers.

### Bottlebrush architecture enables rapid translocation of PEG-based nanocarriers across human airway surface barriers

We use the classic air-liquid-interface (ALI) culture system to model human airway surface barriers^43^. In the ALI system, HBECs are cultured on a porous plastic membrane, through which nutrients are transported from the cell culture medium on the basal side, whereas on the apical side, cells are in contact with air^44^. After approximately 4 weeks, primary HBECs, or human airway epithelial basal cells, differentiate into ciliated cells and goblet cells, forming a pseudostratified columnar epithelium that recapitulates essential biological features of the human airway epithelium^45–48^. Specifically, the pseudostratified airway epithelium consists of three layers: (i) an intact endogenous mucus hydrogel layer, (ii) a periciliary layer that separates the mucus hydrogel from the epithelial cells, and (iii) a layer of epithelial cells connected by cell junctions (**Fig. 2A**)^49^.

**Figure 2.**
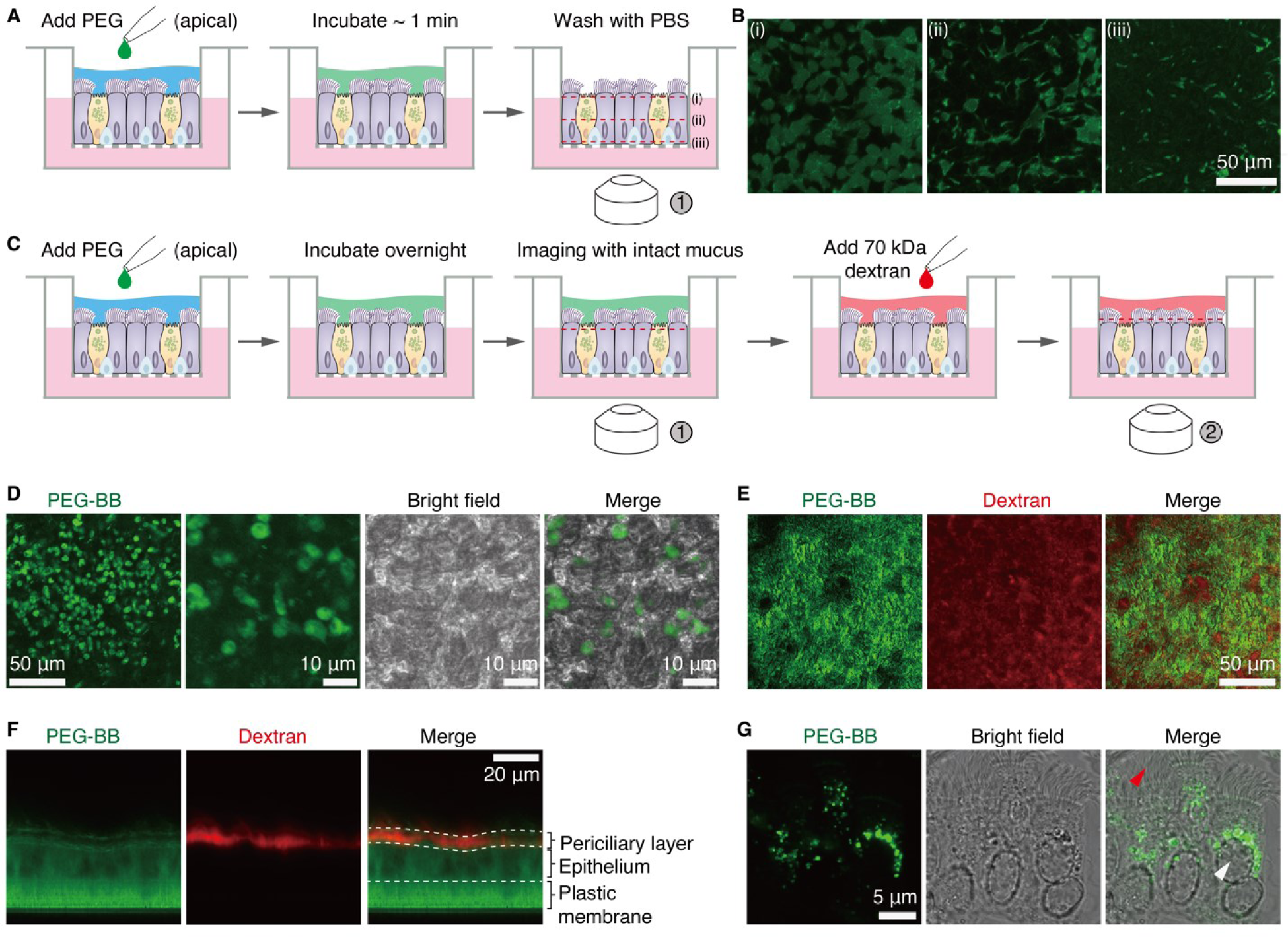
Uptake of PEG-BB by well-differentiated HBECs across the mucus barrier and the periciliary brush layer from the apical side. (A, B) Immediate uptake of PEG-BB by HBECs. **(A)** In this study, 10 μl of 1 mg/ml PEG-BB is added to the apical side of a well-differentiated HBEC culture. After incubation for approximately 1 min, PEG-BB is washed away using DPBS, and then the cells are imaged at different depths into the HBEC layer. **(B)** Fluorescence confocal images showing the uptake of PEG-BB of cells (i) on the apical side, (ii) in the middle, and (iii) on the basal side of the HBEC layer (red dashed lines at timepoint ① in **(A)**). **(C-G)** Long-term uptake of PEG-BB by HBECs with intact mucus. **(C)** In this study, 10 μl of 1 mg/ml PEG-BB is added to the apical side to incubate HBECs overnight. Cells are imaged with intact mucus on the next day. 70 kDa Texas Red^TM^ dextran is added to the apical side to mark the periciliary brush layer and to outline the upper boundary of the epithelium. Cells are imaged both along XY and XZ cross-sections. **(D)** XY images of the cells on the apical side (red dashed line at timepoint ① in **(C)**. **(E)** XY images of the epithelial layer taken at the level of the periciliary brush layer (red dashed line at timepoint ② in **(C)**). **(F)** XZ profile of the epithelial layer, taken at timepoint ② in **(C)**, showing the uptake of 1 MDa PEG-BB (green) but not 70 kDa dextran (red) by HBECs across the whole epithelial barrier. **(G)** PEG-BB accumulates in the cytoplasm but not the nuclei of HBECs. Red arrowhead: cilia; white arrowhead: nucleus.

We start with exploring the uptake of PEG-BB molecules by HBECs from the apical side, where both the mucus hydrogel and the periciliary brush are present to serve as barriers to the delivery of drugs via inhalation^50^. Based on our previous study^6,7^, for well-differentiated HBEC cultures, we allow the mucus to accumulate for two weeks, at which timepoint the mucus reaches a concentration of ∼14% (solids) and a height of ∼15 μm; this corresponds to approximately 1.5 μl mucus per well. To each well we add 10 μl of 1 mg/ml PEG-BB from the apical side so that the final mucus concentration is approximately 2% (solids), comparable to that of healthy mucus. We incubate the culture for about 1 minute and wash off any remaining PEG-BB using pre-warmed Dulbecco’s phosphate-buffered saline (DPBS), as illustrated by **Fig. 2A**. Using fluorescence confocal microscopy, we then immediately image the profile of PEG-BB across the whole epithelial layer. Within such a short period of incubation, HBECs at the apical focal plane exhibit pronounced fluorescence (**Fig. 2B, i**), indicating rapid uptake of PEG-BB by HBECs. These results suggest that PEG-BB molecules can easily penetrate through the mucus and periciliary gels to be internalized by HBECs.

Interestingly, for the cells that contain PEG-BB molecules, the distribution of PEG-BB molecules within individual cells dramatically changes with the depth within the epithelial layer. At the apical focal plane, PEG-BB molecules spread the cross-area of the whole cell, as reflected by the nearly homogenous fluorescence bounded within the contour of the cell cross-section (dashed line, **Fig. 2B,i**). However, as the focal plane moves from the apical to the basal side, the fluorescence area within individual cells dramatically decreases, as shown in **Fig. 2B** and **Movie S1**. This is likely because the incubation time of ∼1 min is too short for PEG-BB molecules to reach the basal side of the epithelial layer.

To explore the cellular uptake over longer durations, after adding PEG-BB to the apical side, we incubate HBECs overnight and then image the distribution of PEG-BB molecules without washing off mucus (**Fig. 2C**). We find that the fluorescence of PEG-BB is heterogeneously present within cells on the apical focal plane, as shown in **Fig. 2D**. To further explore the distribution of PEG-BB across the epithelial layer, we add 70 kDa Texas Red^TM^ dextran to the apical side. As established in our previous study^6^, these molecules penetrate the periciliary layer but cannot cross the epithelial layer, allowing us to delineate the boundary of the epithelial surface, as confirmed by a red fluorescence layer in the middle panel of **Fig. 2E**. Within the periciliary layer, cilia exhibit green fluorescence, suggesting that PEG-BB molecules can accumulate within and fluorescently label the cilia (**Fig. 2E**). Within the epithelial layer, PEG-BB molecules are nearly homogenously distributed, as shown by the bright green fluorescence across the whole epithelium (**Fig. 2F**). By dissociating the pseudostratified epithelium from the Transwell plastic membrane to individual cells, we confirm that PEG-BB molecules are internalized by HBECs; however, PEG-BB molecules are not present in the nucleus and only present in the cytoplasm (**Fig. 2G, Movie S2)**. Further, the side view of the epithelial layer reveals that PEG-BB is within epithelial cells from the apical to the basal side (**Movie S3**). These results demonstrate the cellular uptake of PEG-BB molecules and their ability to penetrate through the whole epithelial layer from the apical side.

Next, we explore the uptake of PEG-BB from the basal side of the airway epithelium, a process critical to the uptake of intravenously administered drugs. We add PEG-BB molecules to the culture medium on the basal side to reach a concentration of 100 μg/ml, incubate the HBEC culture overnight, and change the medium to remove any free PEG-BB (**Fig. 3A**). We find that the cells at the apical focal plane exhibit bright fluorescence (**Fig. 3B**). Moreover, PEG-BB molecules are present within the whole epithelial layer (**Fig. 3C**). Interestingly, compared to the uptake from the apical side, for the uptake from the basal side, although PEG-BB molecules are less abundant within the epithelial cells, they accumulate within the cilia (**Fig. 3C**). This accumulation is further supported by the observation that cilia are also visible due to the presence of PEG-BB (**Fig. 3D**). Nevertheless, these results demonstrate the uptake of PEG-BB by HBECs from the basal side to the apical side.

**Figure 3.**
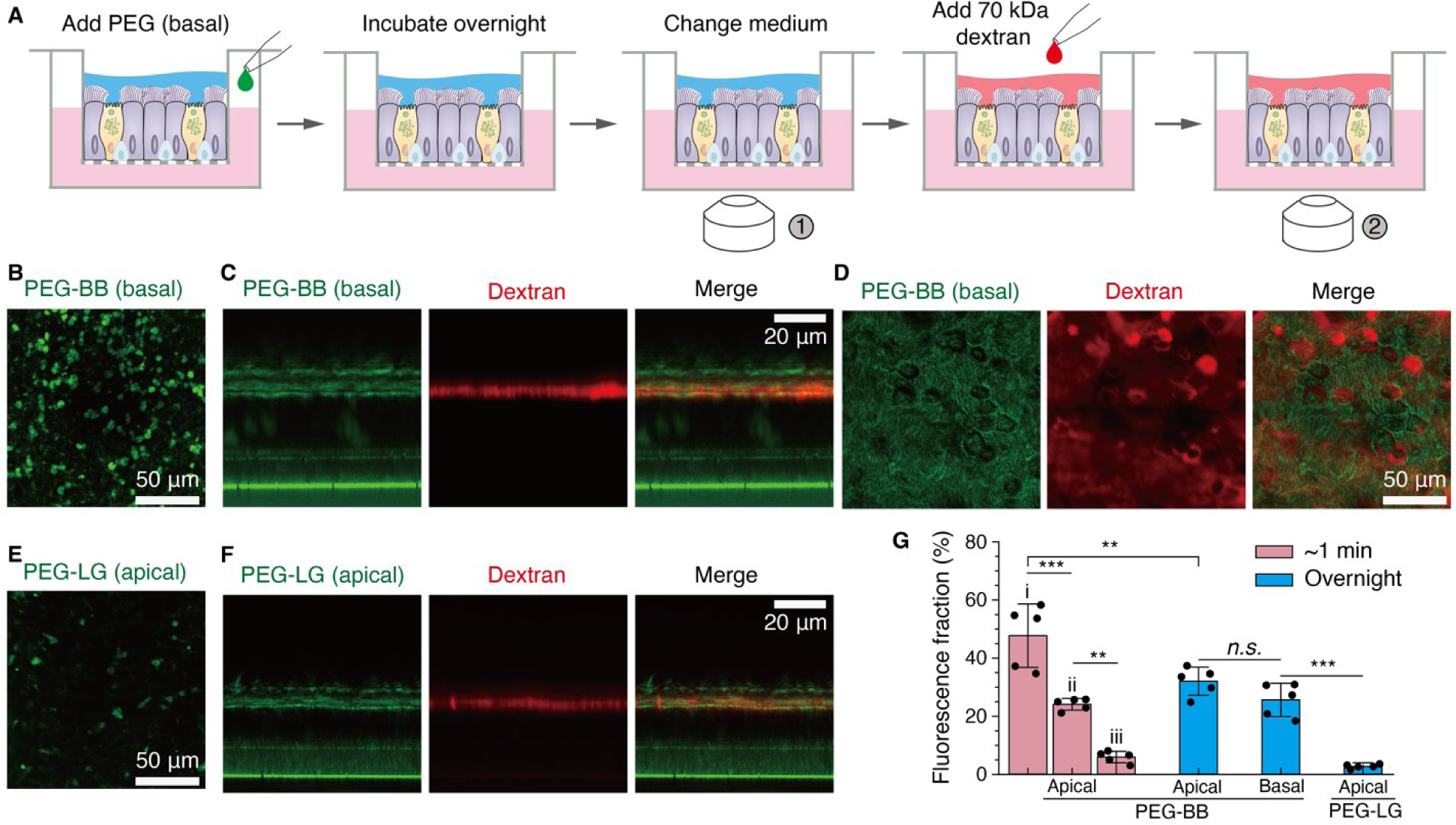
Uptake of PEG-BB from the basal side and PEG-LG from the apical side by HBECs. (a-d) Uptake of PEG-BB by HBECs from the basal side. **(A)** PEG-BB is added to the basal side at the concentration of 100 μg/ml to incubate HBECs overnight. On the next day, the culture medium is changed to remove free PEG-BB, and imaging is performed at the apical focal plane. 70 kDa Texas Red^TM^ dextran is added to the apical side to outline the periciliary brush layer. Imaging of the cells is performed along both XY and XZ profiles. **(b)** A representative XY fluorescence image of HBECs near the apical side (red dashed line at timepoint ① in **(A)**). **(C)** XZ profile of the epithelial layer. **(D)** XY image of HBECs at the level of the periciliary layer (red dashed line at timepoint ② in (A)). **(E, F)** Uptake of PEG-LG by HBECs from the apical side. 10 μl of 1 mg/ml PEG-LG is added to the apical chamber. 70 kDa Texas Red^TM^ dextran is added to the apical side to outline the periciliary brush layer. Imaging is performed at the apical focal plane along both XY and XZ profiles. **(E)** A representative XY fluorescence image of HBECs on the apical side. **(F)** XZ profile of the distribution of PEG-LG in the HBEC epithelial layer. **(G)** Quantitative comparison for the uptake of PEG-BB and PEG-LG by HBECs from the basal and apical sides. Statistical analysis is performed using one-way ANOVA. *n.s.*, not significant; **, *p*<0.01; ***, *p*<0.001. *n=*5 donors.

To determine whether the uptake of PEG-BB molecules is because of their unique bottlebrush molecular architecture, we synthesize a loosely grafted PEG (PEG-LG) polymer with two neighboring PEG macromonomers separated by five spacer monomers on average ([750, 5, 990]), as illustrated by the lower panel of **Fig. 1D** and confirmed by ^1^H-NMR **(Fig. S3)** and GPC (dashed line, **Fig. 1F**). The number of side chains (*n*_*sc*_ = 750) is less than that of PEG-BB (*n*_*sc*_ = 990) to compensate for the contribution of MW by the spacer. Despite that the contour length of PEG-LG (∼1000 nm) is nearly four times of PEG-BB, the grafting density is relatively low at 0.65 nm^-1^, so that the side chains are far apart enough not to experience molecular crowding. As a result, the conformation of PEG-LG is coil-like with a hydrodynamic diameter of 31 nm, comparable to that of PEG-BB (dashed red line, **Fig. 1E**, **Supporting Text**). Similar to PEG-BB, the size of PEG- LG is nearly independent of temperature (dashed red line in **Fig. 1F**). Following the same protocol for studying cellular uptake of PEG-BB, we quantify the uptake of PEG-LG by HBECs from the apical side and find that PEG-LG fluorescence on the apical focal plane is notably reduced (**Fig. 3E**). Profiling the penetration of PEG-LG across the whole epithelial layer further confirms minimum uptake of PEG-LG molecules, as shown by the negligible green fluorescence within the epithelial cells in **Fig. 3F**.

To quantitatively compare the uptake of PEG-BB and PEG-LG by HBECs, we introduce fluorescence fraction, a parameter that is defined as the PEG fluorescence area divided by the total cell area, which is equal to the image area as the cells are confluent. For the immediate uptake of PEG-BB within ∼1 min from the apical side, the fluorescence fraction decreases from 48±11% to 6±2% as the focal plane moves from the apical to the basal side (light red bars, **Fig. 3G**). For overnight uptake, no matter PEG-BB molecules are added from the apical side or from the basal side, there is no significant difference in fluorescence fraction at the apical focal plane (blue bars, **Fig. 3G**). However, changing the molecular architecture from PEG-BB to PEG-LG results in a significant reduction of fluorescence fraction from 32±5% to 3±1% (**Fig. 3G**). These results demonstrate that the bottlebrush architecture significantly enhances the uptake of PEG-based carriers by HBECs.

### Retention of PEG-BB molecules within cells

To explore the retention of PEG-BB molecules within cells, we quantify the cellular uptake of PEG-BB by single NIH-3T3 fibroblasts, a widely used cell line that allows for easy fluorescence staining and manipulation of biological pathways. We add PEG-BB into the culture medium to reach a concentration of 100 μg/ml, incubate NIH-3T3 cells overnight, and replace the medium to wash off any free PEG-BB molecules. Simultaneously, we use Hoechst 33342 to stain the genomic DNA, or nuclei, of the cells. On Day 1, we find that all NIH-3T3 fibroblasts are fluorescent regardless of variations in the area and intensity of fluorescence among individual cells (**Fig. 4A**).

**Figure 4.**
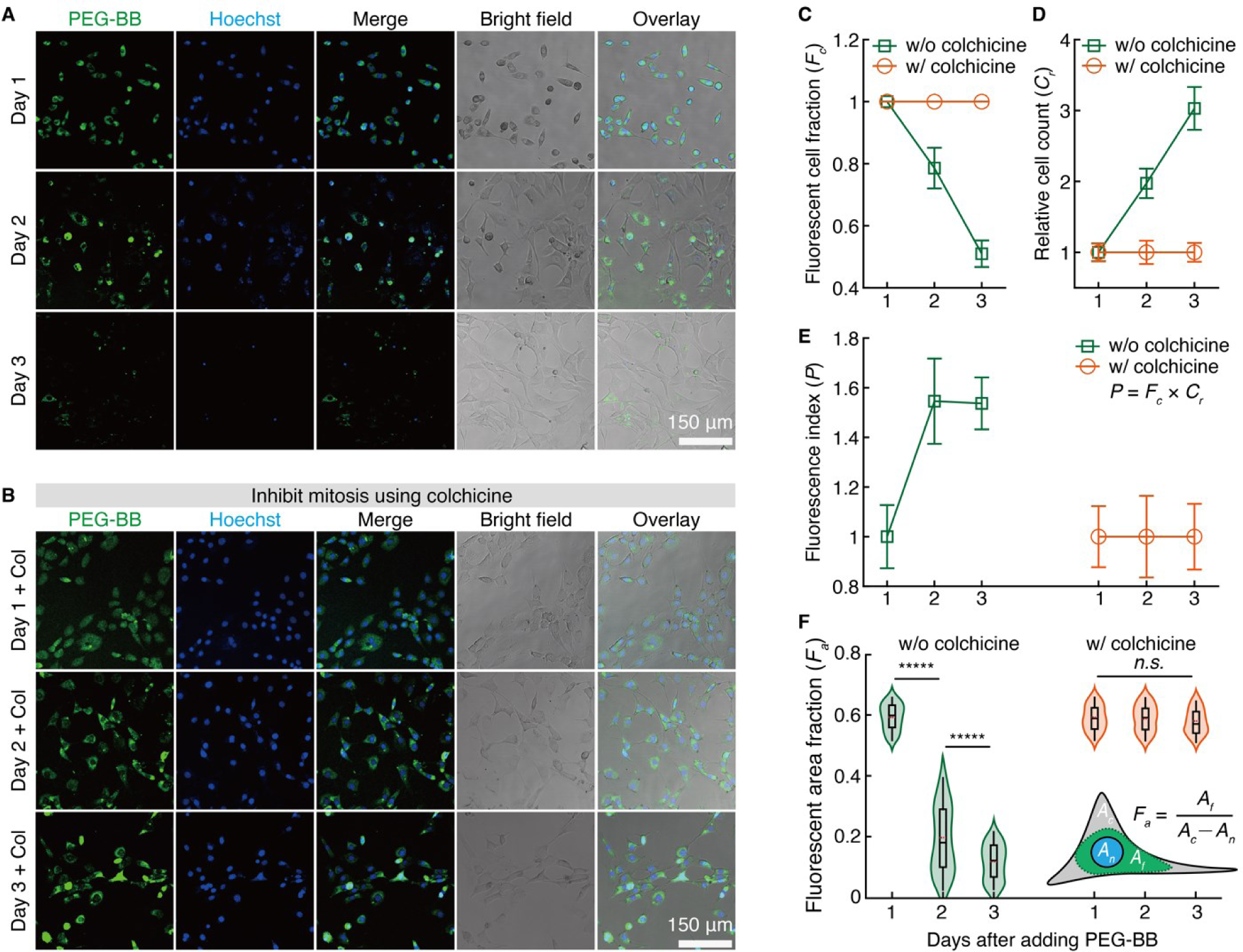
Uptake and retention of PEG-BB molecules within single NIH-3T3 fibroblasts. (A) Uptake of PEG-BB by NIH-3T3 fibroblasts after overnight incubation with 100 μg/mL PEG-BB in the medium on Days 1, 2, and 3. **(B)** Images showing intracellular PEG-BB fluorescence after inhibiting mitosis by 20 ng/mL colchicine, followed by overnight incubation with 100 μg/ml PEG-BB in the medium. Cell nuclei are stained with 20 μg/mL Hoechst 33342 for 5 min at 37 ℃. **(C)** Fluorescent cell fraction (*F_c_*) with and without colchicine treatment. *F_c_* is calculated by the number of fluorescent cells divided by the total cell count. Cells with any fluorescence regardless of the fluorescence area or intensity are counted. *n*=5 wells. **(D)** Relative cell count (*C_r_*) with and without colchicine treatment. *C_r_* is defined as the ratio of cell counts normalized to the initial cell number on Day 1. *n*=5 wells. These results show that the concentration of colchicine is adequate to inhibit mitosis without compromising cell viability. **(E)** Fluorescence index (*P*) with or without colchicine treatment. *P* is defined as the product of *F_c_* and *C_r_*: *P*=*F_c_***×***C_r_. n*=5 wells. **(F)** PEG-BB fluorescent area fraction (*F_a_*) with and without colchicine treatment. *F_a_* is derived by the fluorescent area (*A_f_*) divided by the cytoplasm area. The cytoplasm area is the cell area (*A_c_*) minus the nuclear area (*A_n_*). The results demonstrate that PEG-BB fluorescence is relatively uniformly distributed in the cytoplasm and unchanged over 3 days after inhibiting mitosis. Statistical analysis is performed using one-way ANOVA. *n.s.*, not significant; *****, *p*<0.00001. *n*=100 cells.

However, the number of fluorescent NIH-3T3 fibroblasts dramatically decreases as the culture time increases from Day 1 to Day 3, as visualized by the fluorescence images in **Fig. 4A**. To quantify the number of fluorescent cells, we introduce fluorescent cell fraction *F_c_*, a parameter that is defined as the ratio of the number of fluorescent cells to the total cell count. Specifically, cells with any fluorescence regardless of the fluorescence area or intensity are counted. As the culture time increases from Day 1 to 3, the value of *F_c_* decreases nearly linearly from 100% to 51±4%, as shown by the green squares in **Fig. 4C**. These results show the uptake of PEG-BB across the cell membrane, but that the retention of PEG-BB within the cells decays with time.

To determine the role of cell proliferation in the reduction of PEG-BB fluorescence, we use colchicine to inhibit cell proliferation. Because colchicine inhibits mitosis by disrupting tubulin polymerization^51^, colchicine is cytotoxic and can be lethal to cells at a high dose^52^. To maintain cell viability while inhibiting mitosis, we use colchicine at a concentration of 20 ng/ml^51^ to treat NIH-3T3 fibroblasts before adding PEG-BB. We use relative cell count, *C_r_*, the cell count normalized to that of Day 1, to quantify the proliferation of NIH-3T3 fibroblasts. Without the treatment of colchicine, *C_r_* doubles on Day 2 and triples on Day 3, as shown by the green squares in **Fig. 4D**. By contrast, with the treatment of colchicine, *C_r_* remains constant at 1 across Days 1, 2, and 3, as shown by the orange circles in **Fig. 4D**. These results validate that colchicine at the concentration of 20 ng/ml is adequate to inhibit mitosis without compromising cell viability. With cell proliferation inhibited by colchicine, all NIH-3T3 fibroblasts retain PEG-BB fluorescence across three days without a noticeable decrease in both fluorescence intensity and area among individual cells, as shown by the fluorescence images in **Fig. 4B** and orange circles in **Fig. 4C**. These results indicate that the reduction of intracellular PEG-BB is due to cell proliferation. This understanding is further supported by DAPI fluorescence of cell nuclei, which decreases progressively without colchicine treatment (**Fig. 4A**) but remains nearly constant after inhibiting cell proliferation (**Fig. 4B**).

To further explore the effects of cell proliferation on the retention of intracellular PEG-BB, we introduce fluorescence index, which is the product of fluorescent cell fraction and relative cell count: *P*=*F_c_***×***C_r_*. This parameter describes the total number of cells with intracellular PEG-BB. As expected, after cell proliferation is inhibited by colchicine, the value of *P* remains constant across three days, as shown by the orange circles in **Fig. 4E**. By contrast, for the cells that proliferate, the value of *P* increases by nearly 1.5 times from Day 1 to Day 2; this suggests that PEG-BB molecules are passed to daughter cells during proliferation. Interestingly, at a longer incubation time on Day 3, despite an apparent decrease in fluorescent intensity within individual cells (images within the lower two rows in **Fig. 4A**), the value of *P* remains nearly the same (squares, **Fig. 4E**).

To better understand the retention of PEG-BB within individual cells, we quantify the variation in intracellular PEG-BB fluorescence among different cells. To do so, we introduce fluorescent area fraction, *F_a_*, which is defined as the fluorescent area of a cell, *A_f_*, divided by the cytoplasm area. The cytoplasm area is calculated by subtracting the nuclear area, *A_n_*, from the cell area, *A_c_*, as illustrated by the inset in **Fig. 4F**. As cells proliferate, *F_a_* not only significantly decreases but also shows a higher extent of variation, as shown by the green violin plots in **Fig. 4F**. Note that compared to Day 2, the variation in *F_a_* is lower on Day 3. This is likely because on Day 3 the maximal value of *F_a_* within individual cells is relatively low so that the range of *F_a_* becomes smaller compared to Day 2. Nonetheless, after inhibiting cell proliferation by colchicine, not only there is no significant difference in *F_a_* values but also the variation in *F_a_* remains nearly the same across three days, as shown by the orange violin plots in **Fig. 4F**. Consistent with this understanding, for well-differentiated HBECs, PEG-BB remains intracellular after 7 days (**Movie S4**). Taken together, our results show that the decay of intracellular PEG-BB fluorescence is due to cell proliferation and that PEG-BB molecules remain intracellular for at least 3 days after inhibiting cell proliferation.

### Cellular uptake of PEG-BB nanocarrier is driven by bottlebrush architecture enhanced endocytosis

To further explore the role of molecular architecture in the cellular uptake of polymeric carriers, we quantify the uptake of FITC-labeled 2 MDa dextran by NIH-3T3 fibroblasts. This dextran has a MW on the same order as PEG-BB but is a randomly branched molecule, a molecular architecture that is qualitatively different from the brush-like PEG-BB. After overnight incubation with 100 μg/ml 2 MDa dextran in the culture medium, we find no intracellular fluorescence among all NIH-3T3 fibroblasts (**Fig. 5A, Fig. S4A**). Further decreasing the dextran molecular weight to 70 kDa results in an unmeasurable increase in cellular uptake (**Fig. 5B, Fig. S4B**); this suggests that the molecular weight of dextran molecules has a negligible effect on their uptake by NIH-3T3 fibroblasts. However, dextran and PEG are of different chemical species, which are known to affect the efficiency of cellular uptake^53–55^. To this end, we quantify the uptake of PEG-LG, which has a comparable MW and a hydrodynamic size to PEG-BB but with loosely grafted PEG side chains. Despite that the intracellular PEG-LG fluorescence is slightly higher than that of 70 kDa dextran, it is dramatically lower than that of PEG-BB, as shown by the fluorescence images in **Fig. 5C** and **Fig. S4C**. Together with the minimum uptake of PEG-LG by HBECs, these results confirm that the bottlebrush architecture enables efficient cellular uptake of PEG-BB polymers.

**Figure 5.**
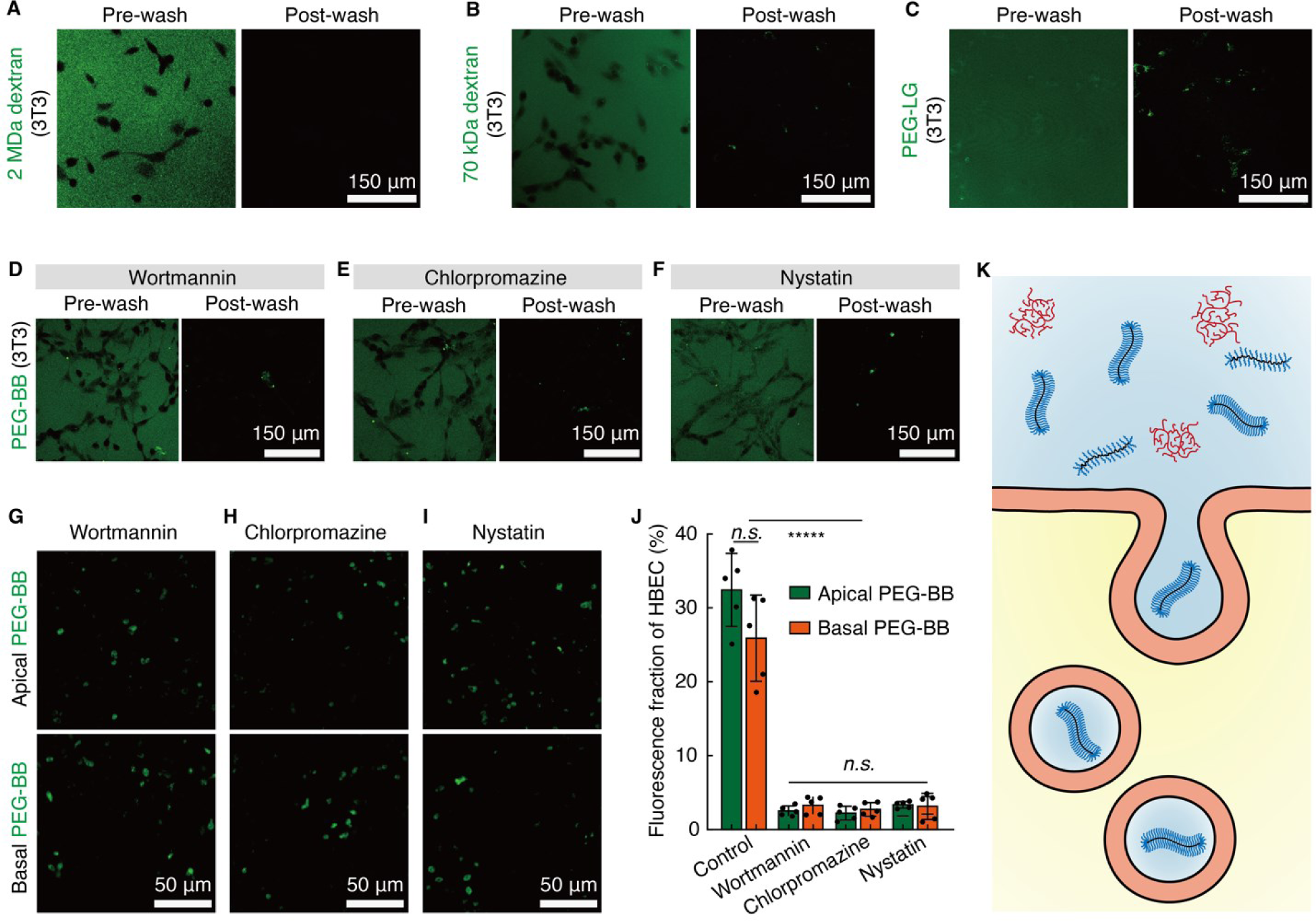
Cellular uptake of PEG-BB is driven by bottlebrush-architecture enhanced endocytosis. (A-C) Uptake of polymers with different molecular architectures by NIH-3T3 fibroblasts: (A) 2 MDa dextran, a randomly branched inert molecule, **(B)** 70 kDa dextran, and **(C)** 1 MDa PEG-LG. All polymers are added to the medium at the same concentration of 100 μg/ml for overnight incubation. **(D-F)** The uptake of PEG-BB by NIH-3T3 fibroblasts is mediated by endocytosis. Representative images of NIH-3T3 fibroblasts treated for 4 hr with **(D)** 5 μg/ml chlorpromazine in the medium, **(E)** 5 μg/ml nystatin in the medium, and **(F)** 0.1 μg/ml wortmannin in the medium. Then, the cells are washed with the prewarmed culture medium and incubated overnight with 100 μg/ml PEG-BB in the culture medium. Pre-wash: cells are imaged right after incubation without replacing the cell culture medium. Post-wash: cells are imaged after being washed with the fresh culture medium. **(G-J)** The uptake of PEG-BB by HBECs is significantly reduced after applying endocytosis inhibitors. Representative images of HBECs after treatment with **(G)** 0.1 μg/ml wortmannin in the medium, **(H)** 5 μg/ml chlorpromazine in the medium, **(I)** 5 μg/ml nystatin in the medium for 4 hr followed by adding 10 μl of 1 mg/ml PEG-BB from the apical side or adding PEG-BB to the basal side achieving a concentration of 100 μg/ml in the medium. Images are for cells right below the apical surface. **(J)** Fluorescence fraction of PEG-BB in HBECs after treatment with endocytosis inhibitors. *n*=5 donors. *n.s.*, not significant; *****, *p*<0.00001. **(K)** A schematic illustrating that the bottlebrush architecture of PEG-BB promotes cellular uptake via endocytosis.

The size of PEG-BB is too large to cross the epithelium by diffusion through cell junctions, which typically necessitate very small molecules of 2∼4 nm or less^56^. Alternatively, endocytosis allows a wide range of substances with various sizes and extents of hydrophobicity to traverse the cell membrane^57^. Given that neither HBECs nor NIH-3T3 fibroblasts are phagocytic cells, we focus on examining whether the cellular uptake of PEG-BB is regulated by pinocytosis pathways. To explore this, we treat cells with wortmannin, a non-specific inhibitor often considered for macropinocytosis which is an endocytosis pathway for nonspecific internalization of large amounts of extracellular fluid^58^. Upon treating NIH-3T3 fibroblasts with wortmannin, nearly all PEG-BB fluorescence is extracellular, outlining the cell contour, as visualized by the left panel in **Fig. 5D** and **Fig. S5A**. Following the removal of free PEG-BB by replacing the medium, negligible fluorescence is detected within NIH-3T3 fibroblasts (right panel in **Fig. 5D** and **Fig. S5A**). Yet, recent studies suggest that wortmannin can also impair clathrin- and caveolin-mediated endocytosis^59^. To identify the specific endocytosis pathways involved in PEG-BB internalization, we treat NIH-3T3 fibroblasts with chlorpromazine which is known to inhibit clathrin-mediated endocytosis^60^ but recently found to impede macropinocytosis^55^. Minimal cellular uptake of PEG- BB is also observed (**Fig. 5E** and **Fig. S5B**). Finally, treating NIH-3T3 fibroblasts with nystatin, an inhibitor of caveolin-mediated endocytosis^61^, effectively prevents the cellular uptake of PEG- BB (**Fig. 5F** and **Fig. S5C**). These results suggest that the internalization of PEG-BB by NIH-3T3 fibroblasts is likely mediated by both macropinocytosis and caveolin-mediated endocytosis.

Based on the knowledge obtained for the uptake of PEG-BB by NIH-3T3 fibroblasts, we apply the three endocytosis inhibitors to well-differentiated HBECs. As expected, all these three inhibitors significantly reduce the cellular uptake of PEG-BB molecules added from both the apical and the basal sides, as shown by the fluorescence images in **Fig. 5G-I**. Quantitatively, for HBECs treated with inhibitors, the fluorescence area decreases from 32±5% to ∼3% and from 26±6% to ∼3% for PEG-BB added to the apical and to the basal sides, respectively (**Fig. 5J**). Yet, the endocytosis pathways that dominate the cellular uptake of PEG-BB remain to be elucidated, which is beyond the scope of this work and will be the subject of future explorations. Nevertheless, our results collectively show that that the internalization of PEG-BB by cells is regulated by bottlebrush architecture enhanced endocytosis (**Fig. 5K**).

## Conclusion and Discussion

We have discovered that bottlebrush PEG polymers, if properly designed, can rapidly penetrate through the mucus gel and the periciliary layer to translocate across the human airway epithelium via molecular architecture enhanced endocytosis. Our PEG-BB is highly anisotropic, featuring a contour length of ∼250 nm, a cross-section of ∼20 nm, and a hydrodynamic diameter of ∼40 nm. The design of PEG-BB draws inspiration from the brushlike mucin biopolymers and mucus-penetrating PEGylated nanoparticles. The size and shape of PEG-BB are based on the mesh sizes of mucus hydrogel (10-100 nm) and periciliary brush layer (20-40 nm). By comparing the cellular uptake of bottlebrush PEG against loosely grafted PEG with a comparable size but a lower grafting density, we show that a high grafting density is critical to efficient cellular uptake. Further, we show that the cellular uptake of PEG-BB is significantly higher than randomly branched dextran molecules regardless of dextran molecular weight. Manipulating the proliferation of NIH- 3T3 fibroblasts reveals that the retention of internalized PEG-BB is determined by cell proliferation rate. Finally, by inhibiting endocytosis pathways, we show the uptake of PEG-BB by fibroblasts and well-differentiated HBECs is regulated by endocytosis.

Compared to existing mucosal delivery systems, the developed PEG-BB serves as a unique nanocarrier in the context of synthesis and design. For example, the ability of the PEGylated nanoparticles to sneak through the sticky mucus hydrogel requires uniform distribution and high grafting density of PEG chains. However, these two parameters are difficult to be precisely controlled because of the nature of the grafting process. Most PEGylated nanoparticles are synthesized using a grafting-through approach, where functional PEG chains are attached to the grafting sites on the surface of the nanoparticle^62^. At a relatively high grafting density, the already highly grafted chains generate steric hindrance to prevent access to the grafting sites^63^. This difficulty is further exacerbated for small nanoparticles (<100 nm) with a relatively high surface curvature^64^. By contrast, PEG-BB is synthesized by polymerizing PEG macromonomers with a prescribed molecular weight (*M*_*sc*_). Within a PEG-BB molecule, the PEG side chains are evenly distributed and have a precisely controlled grafting density (1/(*s* + 1)). Moreover, the total molecular weight of PEG-BB, or the number of side chains (*n*_*sc*_), can be tuned in a wide range through well-established living polymerization techniques. Thus, compared with the PEGylated nanoparticles, the synthesis of PEG-BB is more controlled, enabling prescribed molecular architecture parameters, [*n*_*sc*_, *s*, *M*_*sc*_].

The molecular architecture parameters offer unprecedented control over the geometry and physical properties of PEG-BB as a nanocarrier. For instance, using longer side chains and/or increasing the grafting density increases the cross-section of the bottlebrush, so that the extent of anisotropy of PEG-BB can be tuned for targeted therapeutic delivery^65–68^. An example is that using a small number (∼30) of relatively long PEG chains (10,000 g/mol) results in a PEG carrier with a spherical geometry, which has been demonstrated to enable efficient *in vivo* delivery of nucleic acid therapeutics such as small interfering RNA (siRNA)^69^ and antisense oligonucleotides^70,71^. By contrast, in our studies, PEG-BB consists of many (∼1000) relatively short PEG chains (1 kDa), exhibiting a highly antistrophic, wormlike geometry. Compared to conventional rigid nanoparticles that can be easily trapped within network meshes^72–75^, PEG-BB is a flexible, wormlike nanocarrier, enabling rapid transport through gels and extracellular matrices via reptation^31^. Moreover, drugs for specific diseases can be loaded to and released by PEG-BB using a chemical approach. For instance, multiple kinds of small molecule drugs can be conjugated to macromonomers via cleavable linkers that activate to release drugs, offering strategies for improving monotherapies and combination therapies for multiple myeloma^76^.

We note that at least two open questions are worth future explorations. First, in the human airway epithelium, what are the types of cells that exhibit the most abundant cellular uptake of PEG-BB? This is critical for identifying respiratory diseases for which PEG-BB is most suitable as a drug carrier. Second, our work is a proof-of-concept study focusing on qualitatively different molecular architectures. It has yet to be systematically determined the effects of all three molecular architectures on the ability of PEG-BB to cross various biological barriers. Nevertheless, together with prescribed molecular design and synthesis, the ability to rapidly penetrate through mucus to be internalized by epithelial cells may pose PEG-BB as a precision nanocarrier^77^ for mucosal drug delivery^78^. Finally, considering that PEG has been used as one of the major polymers for biomaterials research^79,80^ and that biophysical and biochemical properties can be encoded into the molecular architecture, PEG-BB may provide a new building block for biomaterials design.

## Methods

### Materials for polymer synthesis

Fluorescein o-acrylate (Flu, 95%), 2-methoxyethyl acrylate (MEA, 98%, monomethyl ether hydroquinone (MEHQ) as inhibitor), and poly(ethylene glycol) methyl ether methacrylate (MEMA-PEG, 950 g/mol, MEHQ as inhibitor) are purchased from Sigma Aldrich and purified by recrystallizing in acetone to remove inhibitors. 2- (dodecylthiocarbonothioylthio)-2-methylpropionic acid (DDMAT, 98%), 2,2’-azobis(2- methylpropionitrile) (AIBN, 98%) and *N, N*-dimethylformamide (DMF, ≥99.8%) are purchased from Sigma Aldrich and used as received.

### Synthesis of PEG-based carriers

<u>I. Densely grafted bottlebrush PEG (PEG-BB).</u> A 25 mL Schlenk flask is charged with MEMA-PEG (2.28g, 1mmol, 1200 eq), Flu (12eq), DDMAT (1 eq), AIBN (0.2 eq), and 6 mL DMF. We de-gas the mixture using three freeze-evacuate-thaw cycles and then seal the flask under nitrogen. We immerse the sealed flask in a heated oil bath at 70 °C for 12 hours and then stop the reaction by exposing the solution to air, at which timepoint the conversion of the polymerization is 82.5%, as confirmed by ^1^H NMR (**Fig. S2**). At this conversion, the DP of PEG is 990, and the DP of Flu is about 10. The reaction mixture is precipitated in ethyl ether three times to remove unreacted monomers and other impurities. Using a dialysis tube with a molecular weight cutoff (MWCO) of 3.5 kDa, we further purify PEG-BB through dialysis against water for 3 days. The solution is freeze-dried for 3 days to obtain the final product, which is a light-yellow powder.
<u>II. Loosely grafted PEG (PEG-LG).</u> A 25 mL Schlenk flask is charged with MEMA-PEG (2.14g, 2.25mmol, 750eq), MEA (1.85g, 11.4mmol, 3800eq), Flu (45.5eq), DDMAT(1eq), AIBN (0.2eq), and 10 mL DMF. We de-gas the mixture through three freeze-evacuate-thaw cycles and then seal the flask under nitrogen. Then, we place the sealed flask in a heated oil bath (70 °C) for 12 hours. The reaction is stopped by exposing the solution to air, with the final DP of PEG is about 750. The reaction mixture is precipitated in ethyl ether and dialysis against water for 3 days using tubes with a pore size molar mass cutoff of 3.5 kDa. Then, we freeze-dry the solution under vacuum for 3 days to get the final product. The final molar ratio between MEMA-PEG and MEA is 1:5, as confirmed by ^1^H NMR (**Fig. S3**).

### Characterization of PEG-based carriers

1. ^1^H NMR characterization. Proton nuclear magnetic resonance (^1^H NMR) spectroscopy is performed using Varian NMRS 600 MHz spectrometer. For all samples, deuterated chloroform (CDCl_3_) is used as a solvent, except for PEG-LG which is analyzed using deuterated water (D_2_O).
2. <u>Gel permeation chromatography (GPC) characterization.</u> GPC measurements are conducted using TOSOH EcoSEC HLC-8320 GPC system equipped with two TOSOH Bioscience TSKgel GMHHR-M 5 µm columns in series. The GPC system includes a refractive index detector and operates at a temperature of 40 °C. High-performance liquid chromatography (HPLC) grade trifluoroacetic acid (TFA) is used as the eluent, and it is delivered at a flow rate of 1 mL/min. The samples for analysis are prepared by dissolving them in TFA at a concentration of approximately 5 mg/mL.
3. <u>Dynamic light scattering (DLS).</u> DLS and ζ-potential measurements are performed on a Malvern Zetasizer Ultra with a 4.0 mW laser (633 nm) at different temperatures. Samples are dissolved in water with a concentration of 0.2 mg/ml and filtered (0.45 μm, PTFE) before measurement. Size measurements are performed in square DTS0012 cuvettes (Malvern) in triplicate. For PEG-BB and PEG-LG, the ζ-potential values are -8.8 mV and -2.0 mV, respectively.

### Human bronchial epithelial cell (HBEC) culture

Primary HBECs are obtained from Marsico Lung Institute Tissue Procurement and Cell Culture Core at the University of North Carolina at Chapel Hill, under protocol number 194 03-1396 approved by the UNC Biomedical Institutional Review Board. For statistics, we use HBECs from 5 non-smoker (NS) donors without a history of chronic lung diseases (age/sex/race: 49/Female/Caucasian, 17/Male/Caucasian, 30/Female/Hispanic, 27/Female/Caucasian, 22/Female/Caucasian). For cell expansion, we culture HBECs using PneumaCult™-Ex Plus Medium (STEMCELL Technologies, Cat. No. 05040). We passage HBECs using Accutase^TM^ Cell Detachment Solution (Innovative Cell Technologies, Cat. No. AT 104). For all experiments, we use passage 2 cells, beyond which HBECs may lose their stemness^44^. We seed HEBCs on Transwell^®^ inserts (Corning, Cat. No. 3460) at the density of 4.2×10^4^ cells/well and add PneumaCult™-Ex Plus Medium (STEMCELL Technologies, Cat. No. 05040) to both the basal and apical chambers. After 5-7 days, HBECs reach over 90% confluence. We transition the cultures to ALI by removing the apical medium and replacing PneumaCult™- Ex Plus Medium with PneumaCult™-ALI Medium (STEMCELL Technologies, Cat. No. 05001). The medium is changed every other day. After 2 weeks of ALI culture, mucus starts to accumulate and is washed three times per week. To wash off mucus, we add 500 μl of Dulbecco’s phosphate-buffered saline (DPBS, Gibco, Cat. No. 14-200-075) to each insert, incubate the cell culture for 15 min, and aspirate the DPBS in the apical chamber. The washing process is repeated three times. After 4 weeks of ALI culture, HBECs are fully differentiated. Based on our previous study^6,7^, we allow the mucus to accumulate for approximately 2 weeks without washing, reaching a concentration of ∼14% solids and a height of ∼15 μm.

### Measurement of the translocation of PEG-based carriers across the airway epithelial layer

To measure the uptake of PEG-BB from the apical side, we add 10 μl of 1 mg/ml PEG-BB to the apical side of each HBEC culture, incubate the culture for about 1 min, and then rinse the culture with pre-warmed DPBS to wash off any remaining PEG-BB. We use fluorescence confocal microscopy (Leica, SP8) to quantify the uptake of fluorescent PEG-BB. The confocal microscope is equipped with an environmental chamber, which has a controlled temperature at 37℃, CO_2_ at 5%, and humidified air at 2L/hr, to allow for long-time live-cell imaging. For fluorescein, a 512 nm laser is used for excitation, and a bandwidth of 500-600 nm is used for emission. Using Z-stack scanning, we image the full thickness of the cell body of the airway epithelium at the step size of 1.12 μm with a total of 15 frames. The periciliary layer is about 7 μm^6^. Therefore, the whole airway epithelium is around 23 μm thick, consistent with the literature value^47^.

To explore the cellular uptake over longer durations, we add 10 μl of 1 mg/ml PEG-BB to the apical side and incubate HBECs overnight. Imaging is performed to show the distribution of PEG-BB molecules without washing off mucus. After performing initial confocal microscopy with intact mucus, we add 10 μl of 100 μg/ml 70 kDa Texas Red^TM^ dextran (Thermo Fisher Scientific, Cat. No. D1830) solution to the apical side and wait for about 1∼2 hours. The 70 kDa dextran molecules penetrate the periciliary layer but cannot cross the epithelial layer, allowing us to delineate the boundary of the epithelial surface^6^. We use confocal microscopy to image the full profile of the culture using XZ scanning mode and to image the fluorescence of the cells at the apical focal plane using XY scanning mode.

To explore the uptake of PEG-BB from the basal side of airway epithelium, we add PEG- BB molecules to the culture medium in the basal chamber to reach a concentration of 100 μg/ml, incubate the HBEC culture overnight, and change the medium on the next day to remove any free PEG-BB. We use confocal microscopy to capture fluorescence at the apical focal plane.

To determine the effect of molecular architecture on the uptake of PEG, we study the absorption of loosely grafted PEG (PEG-LG) by HBECs from the apical side. We add 10 μl of 1 mg/ml PEG-LG to the apical side of HBECs overnight incubation. Imaging is performed to show the distribution of PEG-LG molecules without washing off mucus. A similar procedure is used to label the periciliary layer followed by confocal imaging.

### Imaging of PEG-BB in HBEC cytoplasm

We detach differentiated airway epithelial cells from the Transwell membrane by incubating HBECs with 1 ml of Accutase™ Cell Detachment Solution per well for 15 min at 37 ℃. We add 200 μl of PneumaCult™-ALI Medium to a rectangle coverslip and use a 1000 μl pipette tip to scrape a full thickness of airway epithelium into the medium. Another rectangle coverslip is used to cover the medium containing the airway epithelial sample. Afterward, the sample is mounted onto a 63× oil objective (NA 1.4) with pre-applied lens oil for confocal microscopy. For fluorescein, a 512 nm laser is used for excitation, and a bandwidth of 500-600 nm is used for emission.

### NIH-3T3 cell culture, imaging of PEG-BB fluorescence, and mitosis inhibition by colchicine

For the culture of NIH-3T3 cells, we use Dulbecco’s Modified Eagle Medium (Corning, Cat. No. 10-013-CV) supplemented with 10% Fetal Bovine Serum (Life Technologies Corporation, Cat. No. A3160401). The medium is changed every other day. We passage cells using Accutase^®^ Cell Detachment Solution (Innovative Cell Technologies, AT 104).

To study the internalization of PEG-BB by NIH-3T3 fibroblasts, we add PEG-BB to the culture medium to reach a concentration of 100 μg/ml and incubate cells overnight. On the next day, the culture medium is replaced with the fresh medium to wash off any free PEG-BB. After washing, we stain cell nuclei by adding Hoechst 33342 (Thermo Fischer Scientific, 62249) to medium at the concentration of 20 μg/ml and incubating NIH-3T3 fibroblasts for 5 min at 37 ℃, followed by rinsing the cells twice with DPBS to wash away remaining Hoechst 33342 molecules. Confocal microscopy is performed to image intracellular PEG-BB fluorescence.

To inhibit mitosis without impairing the viability of NIH-3T3 fibroblasts, we choose 20 ng/ml colchicine (Thermo Fisher Scientific, Cat. No. 227120010) to treat NIH-3T3 fibroblasts for 30 minutes at 37 ℃ before treatment with PEG-BB^51^. We use the same protocols as above to incubate NIH-3T3 fibroblast with PEG-BB, stain nuclei, and image intracellular PEG-BB fluorescence.

For each NIH-3T3 culture, we use a 10× dry objective to image a region of interest (ROI) with a dimension of 381.5×381.5 μm^2^. The sequential scanning mode is used to avoid overlap of fluorescence. For fluorescein, a 512 nm laser is used for excitation, and a bandwidth of 500-600 nm is used for emission. For Hoechst 33342, a UV laser of 405 nm is used for excitation and a bandwidth of 400-500 nm is used for emission.

### Measurement of the cell number, fluorescent cell fraction, fluorescence index, and fluorescent area fraction

We count the cell number based on the number of nuclei stained by Hoechst 33342. We use relative cell count (*C_r_*), defined as the ratio of cell counts normalized to the initial cell number on Day 1, to present the cell number on each day.

Fluorescent cell fraction (*F_c_*) is defined as the ratio of fluorescent cells to the total cell count. Regardless of the size or intensity of fluorescence, cells with any fluorescence are counted. For each condition, 5 parallel wells of cell culture are used for statistical analysis. Fluorescence index (*P*) is defined as *P* = *F_c_*×*C_r_*.

Fluorescent area fraction (*F_a_*) is derived by the fluorescent area (*A_f_*) divided by the cytoplasm area. The cytoplasm area is the cell area (*A_c_*) minus the nuclear area (*A_n_*). For each condition and timepoint, 100 cells are randomly chosen for statistical analysis.

### Uptake of PEG-BB, 2 MDa dextran, 70 kDa dextran, and PEG-LG by NIH-3T3 fibroblast cells

PEG-BB, 2 MDa FITC dextran (Millipore Sigma, Cat. No. FD2000S), 70 kDa FITC dextran (Millipore Sigma, Cat. No. FD70), and PEG-LG are added to the culture medium at the same concentration of 100 μg/ml to incubate with NIH-3T3 cells overnight respectively. After incubation, we use 20 μg/ml Hoechst 33342 to incubate NIH-3T3 cells for 5 min at 37 ℃ to stain the cell nuclei and then DPBS to rinse the culture twice. After rinsing, the same amount of 2 MDa FITC dextran, 70 kDa FITC dextran, and PEG-LG are re-added for pre-washing imaging using confocal microscopy. Washing with pre-warmed medium is performed twice, followed by post-washing imaging using confocal microscopy.

### Effects of endocytosis inhibitors on the uptake of PEG-BB by NIH-3T3 fibroblasts and HBECs

We add 5 μg/ml chlorpromazine (Thermo Fisher Scientific, Cat. No. J63659), 5 μg/ml nystatin (Thermo Fisher Scientific, Cat. No. BP29495), and 0.1 μg/ml wortmannin (Thermo Fisher Scientific, Cat. No. W0007) to the culture medium to incubate NIH-3T3 fibroblasts and HBECs for 4 hours at 37 ℃, to inhibit endocytosis. After inhibiting endocytosis, we study the cellular uptake of PEG-BB by NIH-3T3 fibroblasts and HBECs using the same protocols described above. To study how inhibition of endocytosis affects PEG-BB uptake by NIH-3T3 cells, we add 100 mg/ml PEG-BB to the culture medium and incubate cells overnight. On the next day, we stain cell nuclei by adding Hoechst 33342 to the culture medium at a concentration of 20 μg/ml and incubating NIH-3T3 cells for 5 min at 37 ℃, followed by rinsing the cells twice with DPBS to wash away free Hoechst 33342. We perform confocal microscopy to capture both extracellular and intracellular PEG-BB fluorescence if any. After initial imaging with the presence of extracellular PEG-BB in the culture medium, we use the fresh medium to wash off any free PEG-BB. After washing, confocal microscopy is performed again to image intracellular PEG-BB fluorescence if any.

To study the uptake of PEG-BB by HBECs from the apical and basal sides, we add 10 μl of 1 mg/ml PEG-BB to the apical side and 100 μg/ml PEG-BB to the basal medium respectively. After overnight incubation, we do not wash the cell culture to keep the mucus layer intact, and we use confocal microscopy to capture fluorescence at the apical focal plane for PEG-BB added from both apical and basal sides.

### Statistical analysis

We perform statistical analysis using one-way analysis of variance (ANOVA). For the post hoc test after performing statistical analysis, we use Tukey’s honestly significant difference (HSD) test to determine the significant differences between groups. *p*>0.05 is considered statistically significant.

## Acknowledgments

We thank Dr. Richard Boucher at UNC Chapel Hill and Dr. Jae-Won Shin at the University of Illinois Chicago for discussions on pulmonary drug delivery and endocytosis pathways. We thank Dr. Kenichi Okuda at UNC Chapel Hill for HBECs, which are provided by the UNC Marsico Lung Institute Tissue Procurement and Cell Culture Core supported by NIH grant DK065988 and CF Foundation grant BOUCHE19R0. L.H.C. acknowledges the support from NSF (CAREER DMR-1944625), NSF CBET-2306012, the University of Virginia (UVA) LaunchPad for Diabetes, the UVA Coulter Center for Translational Research, Juvenile Diabetes Research Foundation (JDRF 1-INO-2022-1114-A-N), grant funding from Virginia’s Commonwealth Health Research Board, and the UVA Center for Advanced Biomanufacturing.

## Author contributions

L.H.C., Z.J.H, and B.H. designed the research. Z.J.H. performed cell studies. B.H. synthesized and characterized PEG-based nanocarriers. L.H.C., Z.J.H., and B.H. wrote the paper. All authors reviewed and commented on the paper. L.H.C. conceived and supervised the study.

## Data availability

All data are available in the manuscript or the supplementary information.

