## Supplemental Information for "Bottlebrush polyethylene glycol nanocarriers translocate across human airway epithelium via molecular architecture enhanced endocytosis"

**This PDF file includes:**

Supporting Text
Figures S1-S5
Movies S1-S4

### Supporting Text

**Molecular structure of a bottlebrush polymer in solution.** In a bottlebrush polymer, the side chains highly overlap with each other, resulting in steric repulsion so that the side chains extend away from the bottlebrush backbone, forming a cylindrical shape with the cross-section illustrated in **Fig. S1A**. To calculate the cross-section of the bottlebrush or the size of a side chain, $R_{sc}$, we consider the profile of the volume fraction, $\phi(r)$, of the side chains at the distance $r$ from the bottlebrush backbone.

$$\phi(r) \approx \frac{b^3 g(r)}{\xi^3(r)}, \qquad \text{for } r > b \qquad \text{(S1)}$$

Here, $b$ is the Kuhn monomer size, $g(r)$ is the number of monomers per correlation blob, $\xi(r) \approx bg^{\nu}(r)$ is the correlation length, and $\nu$ is the Flory exponent depending on solvent quality (for theta solvent $\nu = 1/2$ and for a good or athermal solvent $\nu = 3/5$)[1].

$$\phi(r) \approx \left[\frac{\xi(r)}{b}\right]^{(1-3\nu)/\nu}, \qquad \text{for } r > b \qquad \text{(S2)}$$

Within the cylinder-like bottlebrush polymer, the correlation length is related to the linear grafting density $1/l$ of side chains[2,3]:

$$\xi(r) \approx (rl)^{1/2}, \qquad \text{for } r > b \qquad \text{(S3)}$$

which increases the distance $r$ by a power of 1/2 (**Fig. S1B**). Note that in our bottlebrush polyethylene glycol (PEG-BB) polymer, the grafting distance is very small with $l = 0.254$ nm, much smaller than the size $b = 0.8$ nm of a polyethylene glycol (PEG) Kuhn monomer[4,5]. Thus, at the length scale $r < b$, the side chains fill the space to form an exclusion zone for solvents, as predicted by the previous theory[6] and confirmed by simulation[7], as well as illustrated by the shadowed circle in **Fig. S1A** and the dashed line in **Fig. S1B**. However, the exclusion zone is very small on the order of Kuhn monomer size. Thus, we ignore the effect of exclusion zone on the bottlebrush thickness and focus on the region with $r > b$, where the polymer chains interact with solvent molecules.

Substituting eq. (S3) to eq. (S2), the volume fraction profile can be re-written in terms of the distance $r$ from the bottlebrush backbone:

$$\phi(r) \approx \left(\frac{rl}{b^2}\right)^{(1-3\nu)/2\nu}, \qquad \text{for } r > b \tag{S4}$$

The size of a side chain, $R_{sc}$, can be determined based on mass conservation:

$$\int_b^{R_{sc}} \phi(r)\, \xi^2(r) dr \approx N_{k,sc} b^3 \tag{S5}$$

where $N_{k,sc}$ is the number of Kuhn monomers per side chain. Solving eq. (S4) one obtains:

$$R_{sc} \approx \left(\frac{b}{l}\right)^{\frac{1-\nu}{2\nu}} b\left(N_{k,sc}\right)^{\frac{2\nu}{1+\nu}} \tag{S6}$$

The bottlebrush polymer is effectively a 'fat' linear polymer with an effective Kuhn length about the cross-section of the bottlebrush, $b_e \approx 2R_{sc}$. The radius gyration of this 'fat' linear polymer is proportional to the end-to-end distance:

$$R_g = \alpha b_e \left(\frac{L_{max}}{b_e}\right)^{\nu} \quad \text{(S7)}$$

where $\alpha = 0.4$ for a linear polymer in good solvent[8] and $L_{max} = n_{sc}l$ is the contour length of the bottlebrush backbone with $n_{sc}$ being the number of side chains per bottlebrush.

Since water is a good solvent for PEG ($\nu = 3/5$), the size of the side chain can be re-written as:

$$R_{sc} \approx \left(\frac{b}{l}\right)^{\frac{1}{4}} bN_{k,sc}^{\frac{3}{4}}, \qquad \text{for good or athermal solvent} \quad \text{(S8)}$$

This suggests that the side chain size not only increases with the grafting density $1/l$ but also scales with the polymer MW by a power of 3/4, higher than 3/5 for an unperturbed linear chain in good solvent.

For PEG in water, the size and mass of a Kuhn monomer are, respectively, $b = 0.8$ nm and $M_0 = 44$ g/mol[4,9]. In our densely grafted PEG-BB, the number of side chains is $n_{sc} = 990$, the grafting distance is $l = 0.254$ nm, and the number of Kuhn monomers per side chain is $N_{k,sc} = M_{sc}/M_0 \approx 22$, in which $M_{sc} = 950$ g/mol is the molecular weight of a PEG side chain. Substituting these numbers into eqs. (S8) and (S7), one obtains the cross-section of the bottlebrush, $b_e \approx 2R_{sc} \approx 20$ nm, and the size of the bottlebrush, $R_g \approx 36\ nm$. The hydrodynamic diameter $d_h$ of a 'fat' linear polymer is related to its radius of gyration $R_g$ by $d_h = 1.25R_g \approx 45$ nm (Table 8.4 in ref. [1]). Considering that scaling theory ignores prefactors on the order of unity, the predicted value agrees reasonably well with the measured hydrodynamic diameter $37 \pm 0.4$ nm (**Fig. 1F**).

For the loosely grafted PEG (PEG-LG), the number of side chains is $n_{sc} = 750$ and the grafting distance $l = 1.524$ nm is much larger than the size of a PEG Kuhn monomer. As a result, the side chains are not much overlapped and adopt a nearly unperturbed conformation. Therefore, the backbone of the bottlebrush polymer is not strained and adopts a self-avoiding random walk with $R_g \approx \alpha b(L_{max}/b)^{3/5} \approx 33$ nm (where $b \approx 1.7$ nm for a methyl methacrylate-based polymer[10] and $L_{max} \approx 1100$ nm is the backbone contour length).

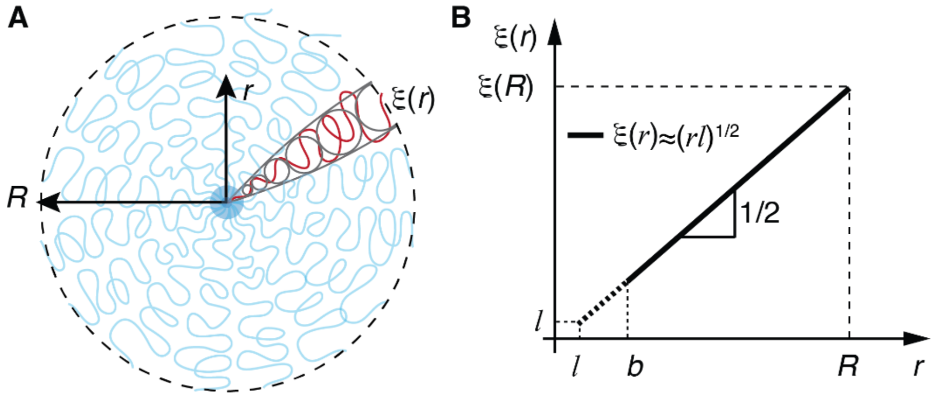


**Figure S1. Cross-section of a bottlebrush polymer in solution. (A)** An unperturbed bottlebrush polymer with the thickness $R$ expected to be larger than the size of the free side chain. The correlation length $\xi(r)$ increases with the distance $r$ from the bottlebrush backbone. **(B)** Mesh size profile for an unperturbed bottlebrush polymer. Yet, at a very high grafting density with the distance $l$ between two neighboring grafting sites much smaller than the Kuhn monomer size $b$, the side chains form an exclusion zone (dashed line), as illustrated by the shadowed light blue circle in **(A)**. Logarithmic scales.

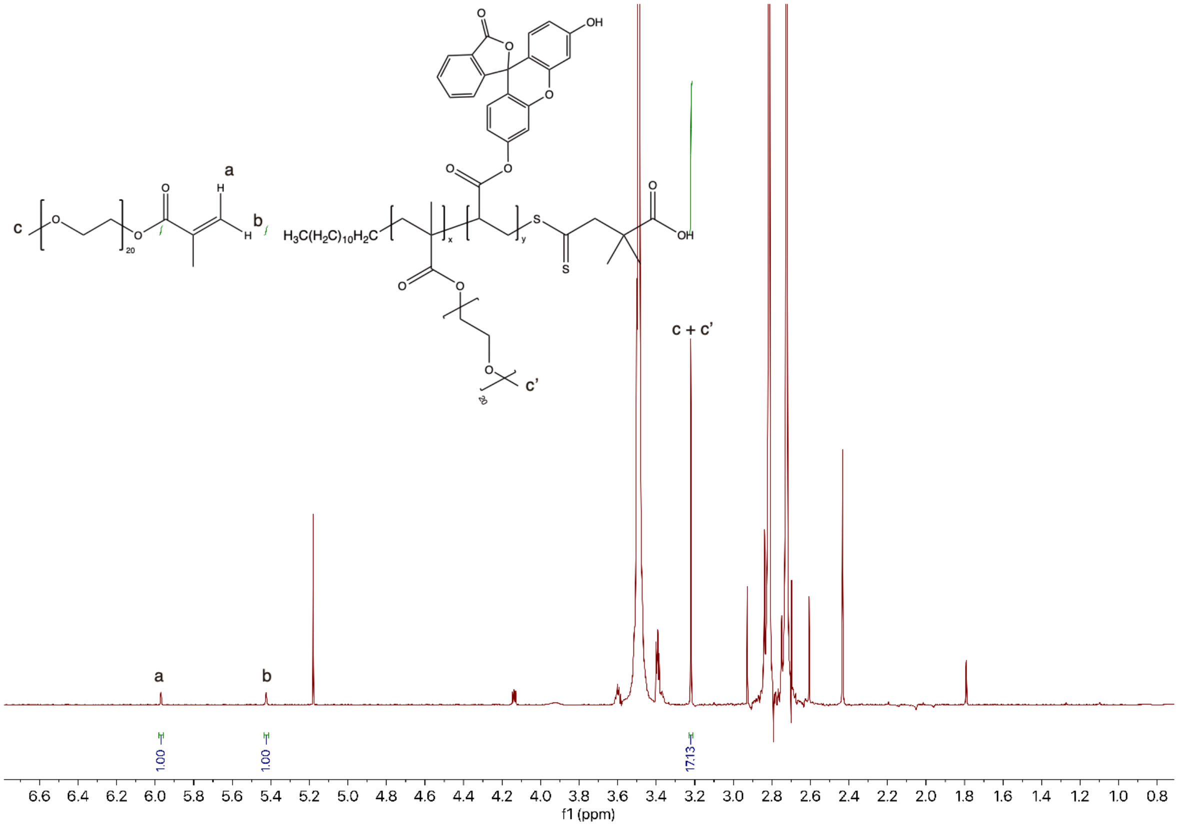


**Figure S2. $^1$H NMR spectra of densely grafted bottlebrush PEG (PEG-BB).** The conversion of PEG is $[1 - (1 \div 17.13) \times 3] \times 100\% = 82.5\%$. The final degree of polymerization (DP) for PEG-BB is $1200 \times 82.5\% = 990$. The feeding molar ratio between PEG and fluorescein *o*-acrylate is 100:1.

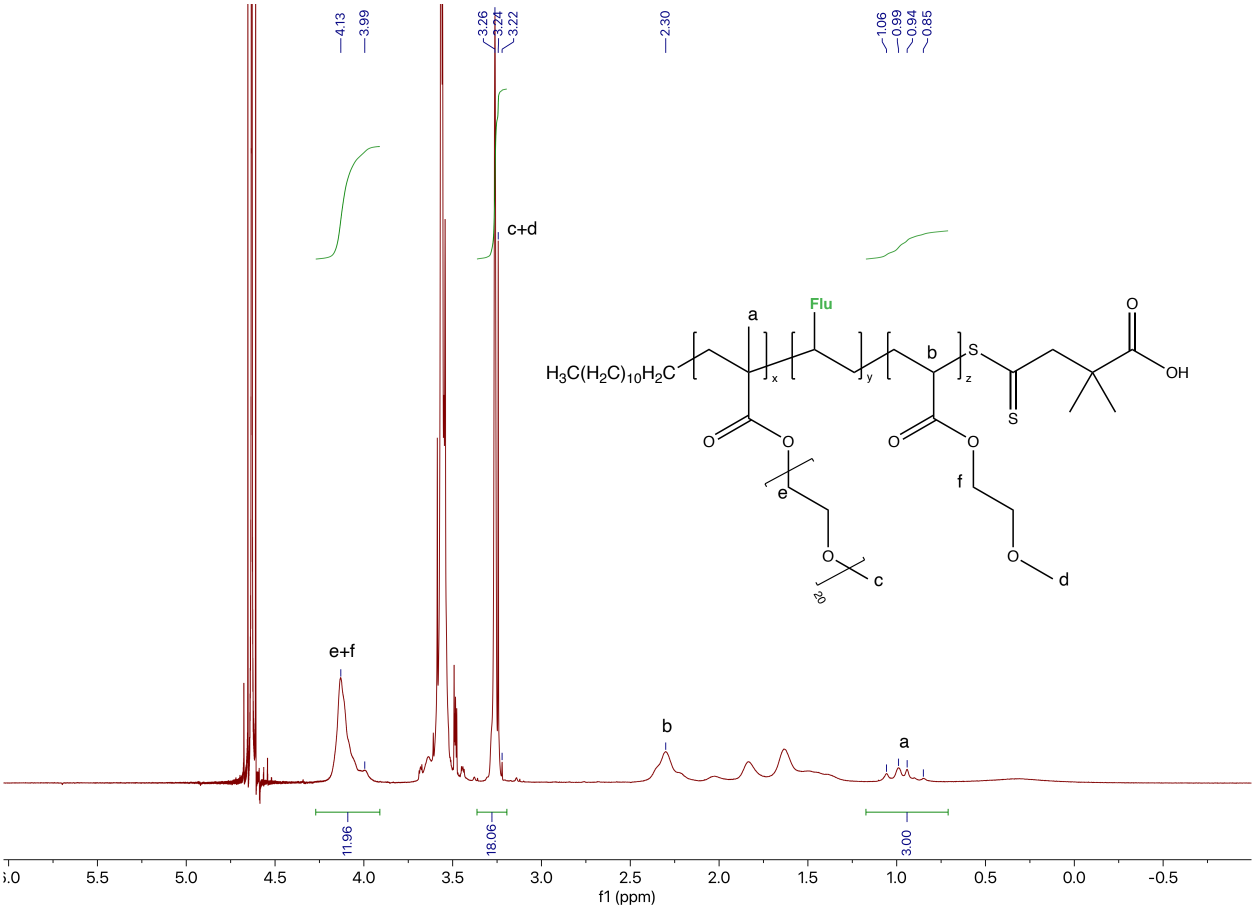


**Figure S3. $^1$H NMR spectra of pure loosely grafted PEG (PEG-LG).** The final molar ratio between PEG and MEA can be calculated in two ways. One is $M_{PEG}$ : $M_{MEA}$ = [Area(e+f)/2-Area(a)/3] : [Area(a)/3], which is [11.96/2-3/3] : [3/3] = 5 : 1. The other way is $M_{PEG}$ : $M_{MEA}$ = [Area(c+d)/3-Area(a)/3] : [Area(a)/3], which is [18.06/3-1]:1 = 5 : 1. The feeding ratio between PEG, MEA, and Flu is 5:1:0.06.

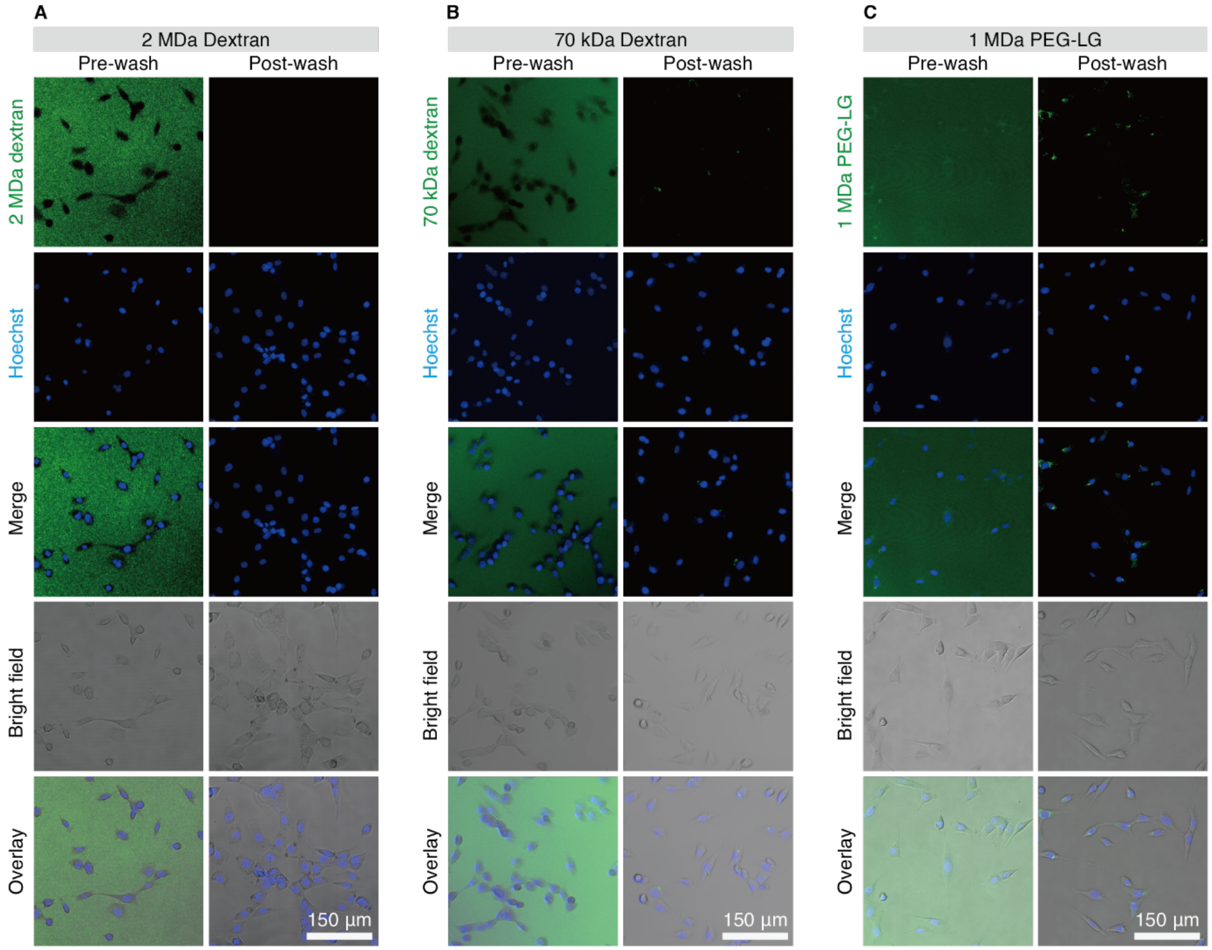


**Figure S4. Uptake of polymers with different molecular architectures by NIH-3T3 fibroblasts.** Representative pre- and post-wash images of NIH-3T3 fibroblasts after overnight incubation with **(A)** 2 MDa dextran, a randomly branched inert molecule, **(B)** 70 kDa dextran, and **(C)** 1 MDa PEG-LG. All polymers are added to the medium at the same concentration of 100 µg/ml. Pre-wash: cells are imaged right after incubation without replacing the cell culture medium. Post-wash: cells are imaged after being washed with the fresh culture medium to remove free extracellular molecules. After overnight incubation, FITC-labeled 2 MDa dextran is not uptaken by NIH-3T3 cells, rendering the cell contour circumscribed. 70 kDa dextran is negligibly uptaken into NIH-3T3 cells as minimal perinuclear green dots. PEG-LG, which has a comparable molecular weight and hydrodynamic size to PEG-BB, is uptaken into NIH-3T3 cells slightly higher than 70 kDa dextran, showing as perinuclear green fluorescence dots. However, the uptake of PEG-LG is significantly less than PEG-BB.

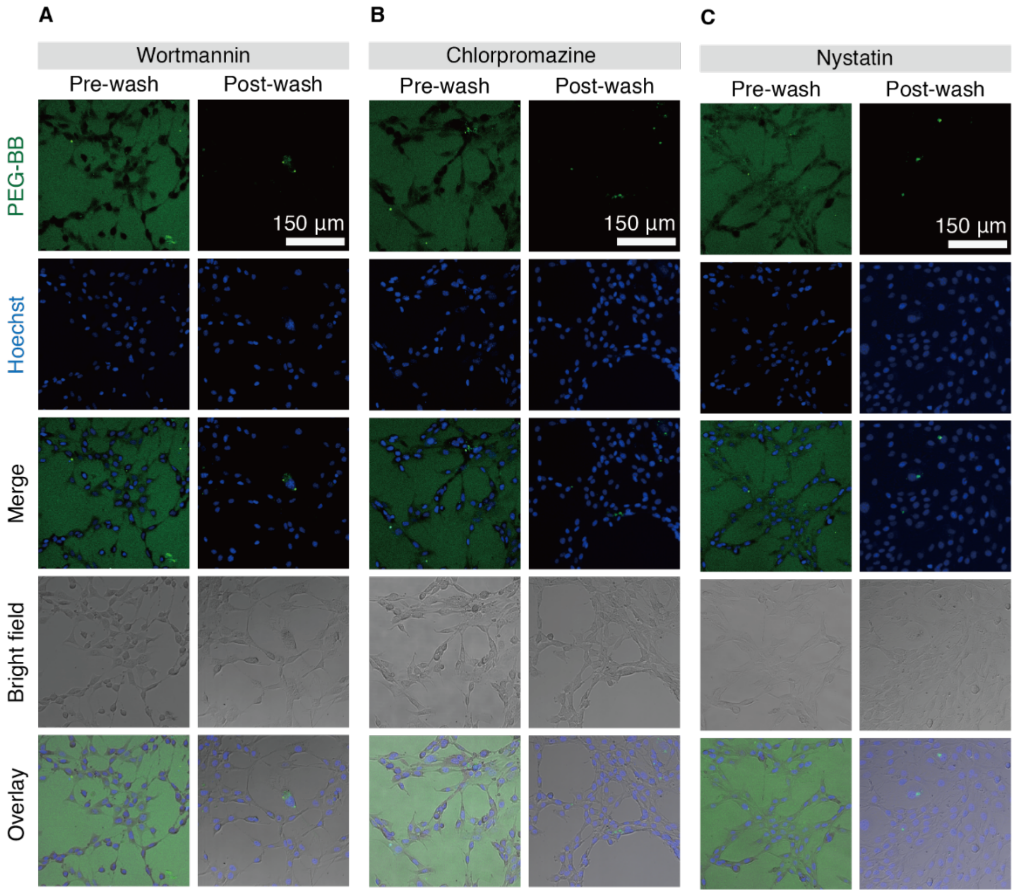


**Figure S5. Cell uptake of PEG-BB is inhibited by endocytosis inhibitors.** Representative images of NIH-3T3 cells treated with **(A)** 0.1 µg/ml wortmannin in the medium, **(B)** 5 µg/ml chlorpromazine in the medium, and **(c)** 5 µg/ml nystatin in the medium for 4 hours. Then, cells are washed with the prewarmed culture medium and incubated overnight with 100 µg/ml PEG-BB in the culture medium. Pre-wash: cells are imaged right after incubation without replacing the culture medium. Post-wash: cells are imaged after being washed with fresh culture medium to remove free extracellular PEG-BB. After being treated with wortmannin, chlorpromazine, and nystatin to inhibit endocytosis, almost all PEG-BB is unabsorbed and remains extracellular, outlining cell contour in the pre-wash images. After washing, minimal PEG-BB remains intracellular and presents as perinuclear green fluorescence dots in the post-wash images.

**Movie S1.** Z profile of PEG-BB uptake by HBECs after adding PEG-BB to the apical side and incubation for ~1 min, followed by rinsing with Dulbecco's phosphate-buffered saline (DPBS). The movie is played at 7 frames per second (FPS). The red dashed line at the upper right corner indicates the imaging focal plane. Images are captured at the step size of 1.12 μm with a total of 15 frames.

**Movie S2.** PEG-BB localizes in the perinuclear cytoplasm of a single ciliated cell 1 day after adding 10 μl of 1 mg/ml PEG-BB to the apical side. Images are captured at 0.346 sec per frame. The movie is played at 15 FPS.

**Movie S3.** The full thickness of airway epithelium contains intracellular PEG-BB 1 day after adding 10 μl of 1 mg/ml PEG-BB to the apical side. Images are captured at 0.690 sec per frame. The movie is played at 15 FPS.

**Movie S4.** HBECs retain intracellular PEG-BB 7 days after adding 10 μl of 1 mg/ml PEG-BB to the apical side. Images are captured at 0.1 sec per frame. The movie is played at 15 FPS.
